# Identification of Antiviral Drug Candidates Against Monkeypox DNA Polymerase and Profilin-like Protein A42R Utilizing an *In-Silico* Approach

**DOI:** 10.1101/2024.08.15.608157

**Authors:** Muhammad Amjid, Muhammad Maroof Khan, Stephen F. Pastore, John B. Vincent, Tahir Muhammad

**Affiliations:** Department of Chemistry, Kohat University of Science and Technology, Kohat Khyber Pakhtunkhwa, Pakistan; Department of Biotechnology, Agriculture University of Peshawar, 25130, Peshawar, Pakistan; Molecular Neuropsychiatry & Development (MiND) Lab, Campbell Family Mental Health Research Institute, Centre for Addiction and Mental Health, Toronto, ON, Canada; Institute of Medical Science, University of Toronto, Toronto, ON, Canada; Department of Psychiatry, University of Toronto, Toronto, ON, Canada

**Keywords:** Monkeypox Virus (MPXV), DNA Polymerase, Profilin-like Protein A42R, Molecular Docking, Molecular Dynamic (MD) Simulation, *In-silico* Screening, Antiviral Compounds

## Abstract

Monkeypox virus (MPXV) is emerging as a major concern in the field of infectious diseases. Current treatments are limited, highlighting the need for new therapeutic options. The use of computational methods, such as molecular docking and molecular dynamic (MD) simulations, is a valuable approach in identifying potential compounds that can target specific proteins of the virus, like the DNA polymerase and profilin-like protein A42R in this case, with the aim of controlling the disease. Our study focused on screening various libraries of compounds for predicted binding to MPXV DPol and A42R proteins, with the top-performing molecules identified based on their docking scores. Among these, Dorsilurin K and Mangostin in complex with DPol, whereas [2-oxo-2-[3-(3,4,5,6-tetrahydro-2H-azepin-7-ylsulfamoyl)anilino]ethyl] 3,5-dimethylbenzoate and N-[4-[2-[4-(4-methylphenyl)sulfonylpiperazin-1-yl]-2-oxoethoxy]phenyl]furan-2-carboxamide in complex with A42R stand out with notably high docking scores, suggesting they may have a good affinity for binding to the DPol and A42R proteins of MPXV respectively. MD simulations confirmed the stability of these ligand-protein complexes followed by evaluation of the ADMET and oral bioavailability analysis. However, it is important that computational methods can suggest promising candidates, *in vitro* and eventually *in vivo* studies are essential to validate these therapeutic candidates. Further studies on these compounds will provide insights into their efficacy, safety, and potential side effects. In conclusion, this study offers promising avenues for developing potential treatments for Monkeypox. If the identified compounds prove effective in further studies, it could be a significant breakthrough in managing this zoonotic disease.

## INTRODUCTION

Monkeypox virus (MPXV), a member of the Poxviridae family, is classified among 11 species, including cowpox ^1^, camelpox, ectromelia ^2^, vaccinia, and variola, and is the causative agent behind monkeypox disease ^1^. The MPXV genome can be divided into three segments: a core region, a left arm, and a right arm. Unlike orthopoxviruses that encode approximately 200 genes, MPXV carries roughly 190 genes. The core region is relatively conserved in the genome and encodes genes related to viral replication and assembly ^3^.

Although less lethal than smallpox, MPXV remains a significant health concern due to its high prevalence and potential for causing disability and disfigurement ^4^. Clinical symptoms include malaise, severe headaches, fever with temperatures ranging from 38.5°C to 40.5°C ^5^, and the development of lesions on the face, extremities, oral mucous membranes, genitalia, conjunctivae, and cornea ^6^. Additional symptoms encompass chills, weakness, fatigue, myalgia, lymphadenopathies in cervical, inguinal, and axillary regions, as well as ulcers and vesicles in the genital and anal areas ^7^. In severe cases, complications such as pneumonia and sepsis can occur, carrying a high risk of mortality ^8^. The incubation period ranges from 7 to 17 days, with fever subsiding three days after the rash onset. Lesions are characterized as painful, indurated, and edematous. Lymphadenopathy, which is absent in smallpox, triggers a robust immune response in monkeypox, distinguishing the two diseases. MPXV has an associated mortality rate of up to 10%, with a particular severity observed in pediatric patients ^4^. MPXV can be transmitted through zoonosis and human-to-human contact. Modes of acquisition include percutaneous exposure, direct contact with skin (especially damaged skin), direct contact with mucous membranes (e.g., oral, vaginal, rectal), and inhalation of infected respiratory particles ^4^. Potential sources of infection encompass infected individuals, animals, or contaminated objects. Notably, direct sexual contact has emerged as a primary mode of transmission in ongoing outbreaks, as indicated by the predominance of anogenital lesions in cases ^9^.

Currently, there are no FDA-approved treatments for monkeypox. However, during the 2022 outbreak emergency, Tecovirimat received FDA approval as the sole antiviral medication to inhibit MPXV replication and manage severe symptoms^5,10^. Additionally, while smallpox medications such as Cidofovir, Brincidofovir, and Tecovirimat have been approved and the smallpox vaccine demonstrates 85% effectiveness against monkeypox ^5,11,12^, it is important to clarify that these treatments also exert their effects against MPXV. Tecovirimat is the sole anti-poxvirus medication officially recommended by the US FDA ^11,12^. While Cidofovir and Brincidofovir inhibit DNA replication and exhibit efficacy against numerous double-stranded DNA viruses^13,14^, Tecovirimat displays a higher specificity for orthopoxviruses, preventing the formation of enveloped virions through inhibition of the conserved protein F13L^15^.

In the present study, we employ an *in-silico* structure-based drug design approach to identify potential lead compounds from six distinct libraries i.e. MedChemExpress natural product library^16^, MPD3 medicinal plant database^17^, alkaloid compound library^18^, MedChemExpress flavonoid compound library^19^, enamine antiviral compound library^20^, and express-pick library^18^ targeting the DPol and A42R proteins of MPXV. We selected DPol and A42R proteins based on the previously reported studies using these to be the most important targets to screen drugs for monkeypox virus. We conducted virtual screening of libraries by performing the molecular docking and subsequently subject the top ten hits to molecular dynamics simulations. These top hit compounds consist of DPol-MPD3_1, DPol - Natural Product_1, DPol-Express Pic_1, DPol-Natural Product_2, DPol-ExpressPic_2, A42R-ExpressPick_1, A42R-ExpressPick_2, A42R-ExpressPick_3, A42R-Emanime_1, and A42R-Natural Product_1. The chemical structures are given in **Figure 1**, and we will discuss these in detail below. Molecular docking and dynamic simulation analyses predict stable binding interactions of these compounds at the DPol and A42R binding sites. Finally, we performed the absorption, distribution, metabolism, excretion and toxicity (ADMET), and oral bioavailibity of the top hit ligands.

**Figure 1:**
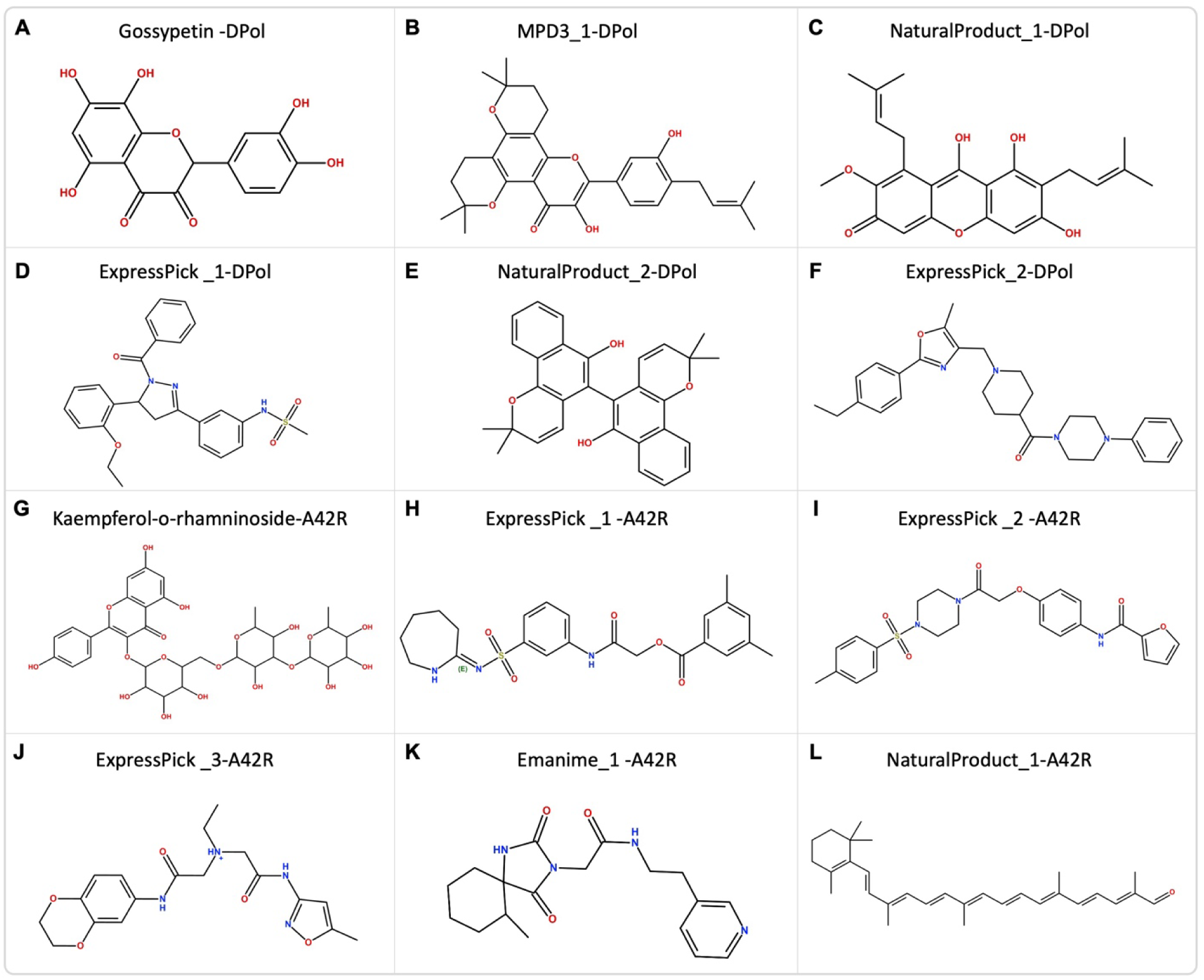
2D chemical Structure of the 10 hit compounds.

## METHODS

### Compound library selection and preparation

Six distinct compound libraries were chosen for analysis, consisting of a natural product library (MedChemExpress), MPD3 medicinal plants database, flavonoid compound library (MedChemExpress), alkaloid compound library (Selleckchem.com), enamine antiviral library, and express-pick library. These libraries comprised 3948, 2295, 241, 401, 3200 and 3010 compounds, respectively, and were obtained in 3D SDF format. To ensure the pharmacokinetic viability of the compounds, an initial filter was applied based on Lipinski’s Rule of Five^21^. After this step, the selected compounds were prepared for subsequent molecular docking processes (**Table 1**).

**Table 1:** Compound libraries, number of total compounds, format and their retrieval date.

| Compound library | Total number of compounds | Selection after Lipinski's rule | Format | Retrieval date |
| --- | --- | --- | --- | --- |
| Natural product library<sup>16</sup> | 3948 | 203 | 3D-SDF | 25 May 2023 |
| MPD3 medicinal plant database<sup>17</sup>8/15/2024 5:37:00 PM | 2295 | 1716 | 3D-SDF | 27 May 2023 |
| Alkaloid compound library<sup>18</sup> | 407 | 314 | 3D-SDF | 14 June 2023 |
| Flavonoid compound library<sup>19</sup> | 241 | 229 | 3D-SDF | 14 June 2023 |
| Enamine antiviral library<sup>20</sup> | 3200 | 3200 | 3D-SDF | 14 June 2023 |
| Express-pick library<sup>18</sup> | 3010 | 2902 | 3D-SDF | 14 June 2023 |

### Protein retrieval and preparation

The X-ray crystal structure of the DPol (PDB ID: 8HG1) and A42R (PDB ID: 4QWO) proteins were retrieved from the Protein Data Bank. The chosen proteins structures underwent further processing using the PyMOL molecular graphics system (Schrödinger, LLC, v2.0)^22^ which involved the removal of redundant chains and DNA molecules. Subsequently, the proteins were imported into the Molecular Operating Environment (MOE) 2022.02 software for preparation^23^. This preparation included steps such as removing water molecules, adding hydrogen atoms, and assigning charges using the OPLS-AA molecular mechanics force field. The active site of DPol and A42R proteins were selected based on the previously reported studies.

### Molecular docking

The site for molecular docking of DPol reported by MA Yousaf et al.^24^ and the MOEs site finder for DPol while the site for A42R reported by Al Mashud et al.^25^ for A42R proteins were designated as potential docking sites. These sites were assigned dummy atoms to facilitate the calculation of molecular docking for the designated sites. The MOE software was then used to perform the ligand-protein docking procedure, utilizing its default parameters.

### Molecular dynamics (MD) simulation

To gain further insights into the stability and interaction dynamics of the top ten docked complexes (two from each library of focus), and MD simulation was initiated. This entire process was carried out using Schrödinger’s Desmond software ^26^. As previously discussed^27^, with few changes, the energy of the entire system was minimized using the OPLS3e force field. A TIP3P water solvent model was applied around the complexes within a 10 Å orthorhombic box. Following ion addition and system neutralization, the system underwent reevaluation. The simulation extended over 100 ns, encompassing 1000 frames, with temperature equilibration set at 300 K and pressure maintained at 1 bar. To ensure system relaxation after minimization, the production phase spanned 20 ns. Output data, including root-mean-square deviation (RMSD), root-mean-square fluctuation (RMSF) and ligand properties were subsequently visualized and analyzed using the Desmond package’s Simulation Interaction Diagram (SID) tool.

### ADMET Analysis

Finally, we performed the ADMET analysis of the selected top hit compounds and compared it to the reported ligands. We first converted the ligands in sdf format to Simplified Molecular Input Line Entry System (SMILE)^28^ followed by ADMET analysis using the swissADME online tool^29^.

## RESULTS

### Molecular docking of DNA polymerase

In the realm of pharmaceutical design, molecular docking studies serve as an invaluable approach for comprehending the interactions between ligands and proteins. Molecular docking, a simulation technique of particular efficacy, employs energy minimization and binding energy calculations to elucidate the interactions between drugs and their target proteins ^30^. Herein, we employed DPol as the target protein, subjecting it to molecular docking with Gossypetin (reference compound) and various other compounds sourced from six distinct libraries, namely the Natural product library, MPD3 medicinal plant database, Alkaloid library, Flavonoid library, Enamine antiviral library, and Express-pick library (**Table 1**). Among these compounds, DPol-MPD3_1 and DPol - Natural Product_1 exhibited the most favorable docking scores, with values of −9.362 and −8.634, respectively (**Table 2**). Additionally, DPol-ExpressPic_1, DPol-Natural Product_2, and DPol-ExpressPic_2, displayed docking scores of −8.640, −8.426, and −8.212 respectively.

**Table 2:** Molecular docking scores, a list of interacting residues, and types of interactions from DPol and A42R proteins of MPXV with the top ten hits.

| Docked complex | Docking scores | H bonds | Ionic bonds | Arene Bonds | Total interactions |
| --- | --- | --- | --- | --- | --- |
| **Gossypetin (Reference) - DPol** | -6.335 | Asp549, Leu631, Arg634, Lys638, Lys638, Glu792 | - | - | 6 |
| MPD3_1 - DPol | -9.362 | Tyr486, Ser655 | - | - | 2 |
| NaturalProduct_1- DPol | -8.634 | Lys337, Ser655 | - | Ser338 | 3 |
| ExpressPick_1- DPol | -8.640 | Ser552, Leu631, Arg634, Lys661 | - | - | 4 |
| MPD3_1- DPol | -8.426 | Ser338 | - | - | 1 |
| ExpressPick_2 - DPol | -8.212 | Ser552, Leu631, Arg634, Lys661 | - | - | 4 |
| **Kaempferol Kaempferol 3-O-rhamninoside (reference) – A42R** | -7.401 | Met1, Met1, Met1, Ala2, Trp4 | - | - | 5 |
| ExpressPick_1- A42R | -7.703 | Glu3, Arg127 | - | His124 | 3 |
| ExpressPick_2- A42R | -7.528 | Met1 | - | - | 1 |
| ExpressPick_3- A42R | -7.382 | Glu3, Arg127 | Glu3 | His124 | 4 |
| Emanime_1- A42R | -7.231 | Met1, His124, Arg127 | - | Trp4 | 4 |
| NaturalProduct_1- A42R | -7.114 | - | - | Trp4 | 1 |

Following the molecular docking analysis, five hits revealed various interactions, including hydrogen bonds and aromatic interactions with DPol (**Figure 2** and **3**, **Table 2)**. DPol-MPD3_1 formed two hydrogen bonds with Tyr486 and Ser655 residues of the receptor, while DPol - Natural Product_1 established two hydrogen bonds with Lys337 and Ser655 residues and one arene bond with Ser338. DPol-Express Pic_1 engaged in four hydrogen bonds with Ser552, Leu631, Arg634, and Lys661 residues, whereas DPol-Natural Product_2 exhibited interaction through hydrogen bond with Ser338 residue. DPol-ExpressPic_2 demonstrated four hydrogen bonds with Ser552, Leu631, Arg634, and Lys661 residues.

**Figure 2:**
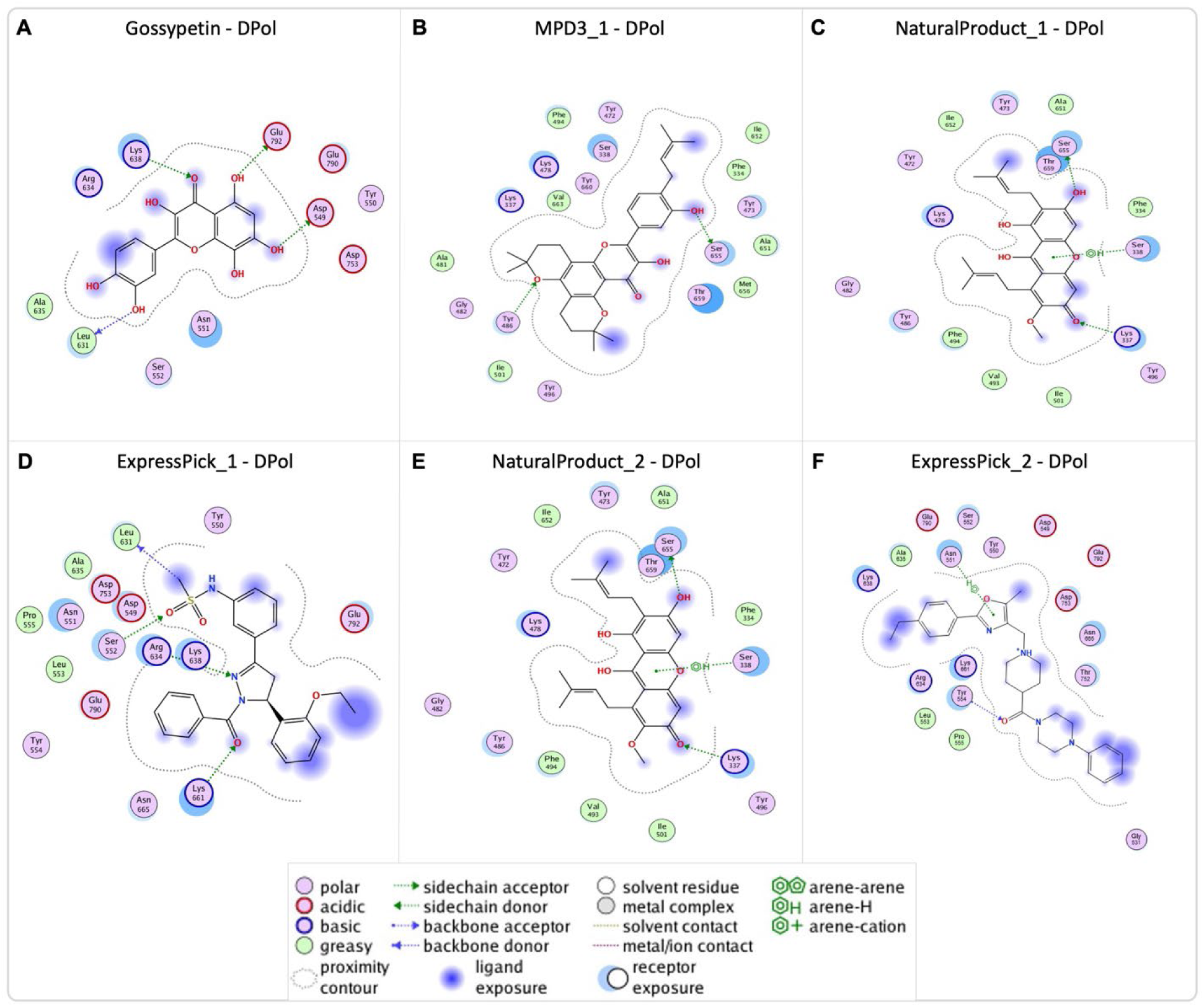
(A-D) 2D structure of the ligands in the binding pocket of DPol, illustrating molecular interactions between each ligand’s atoms and amino acid residues. The types of interactions (hydrogen, ionic, arene) and details regarding ligand exposure, acceptor/donor, polarity, acidic/basic, and greasy or neutral residues of the ligand are presented.

**Figure 3:**
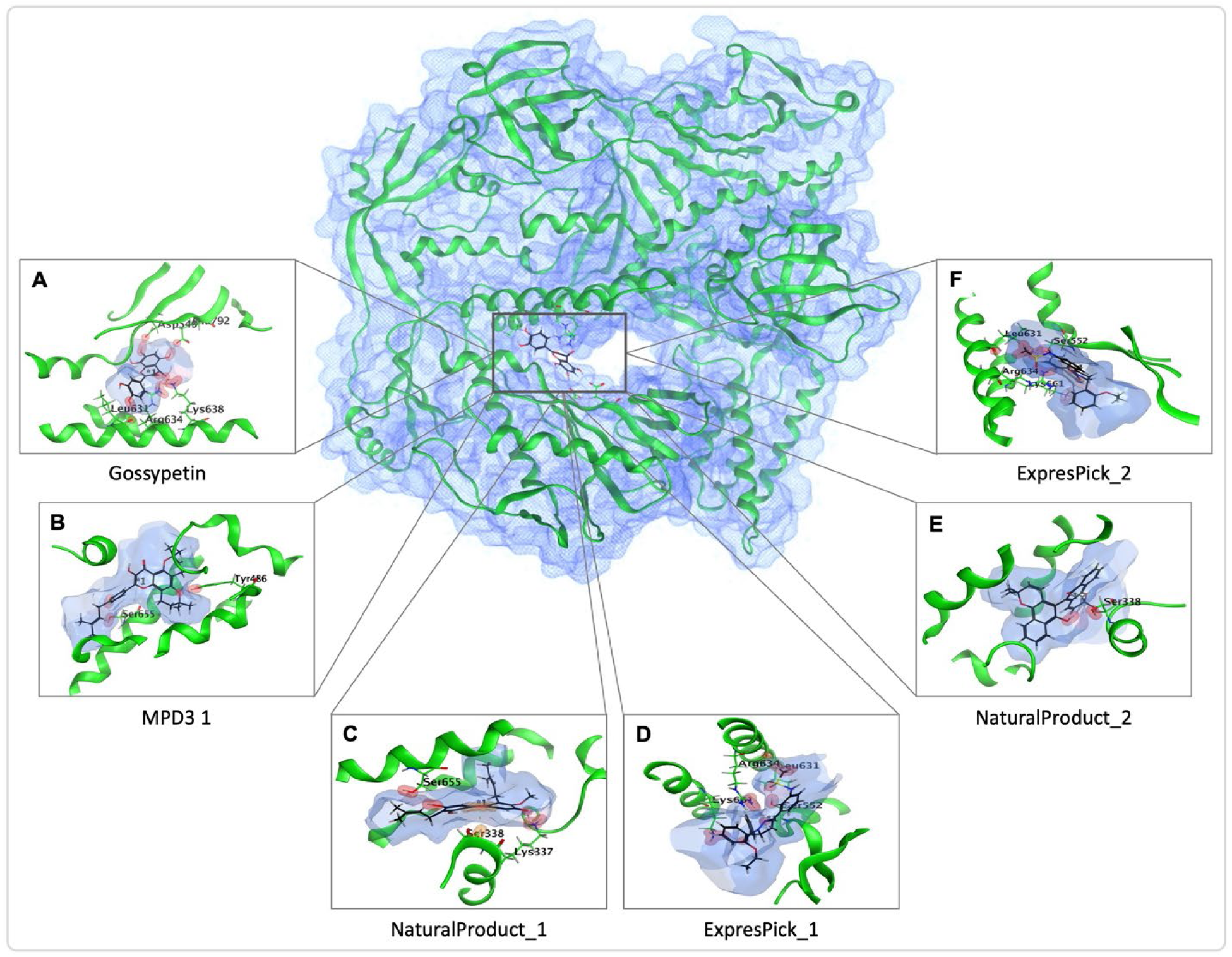
(A-D) 3D representation of the ligands in the binding pocket of DPol, depicting the three-dimensional orientation of the ligands, the names of interacting protein residues, and the ligand’s atoms.

### Molecular docking of profilin-like protein A42R

After performing molecular docking of A42R protein of MPXV with six different compounds libraries, along with Kaempferol 3-O-rhamninoside (reference compound), many compounds showed significant docking scores **(Table 2)**. Among these compounds, A42R-ExpressPick_1 and A42R-ExpressPick_2 have the lowest docking scores, of values −7.703, and −7.528 respectively. In addition to these A42R-ExpressPick_3, A42R-Emanime_1, and A42R-Natural Product_1 displayed docking scores of −7.382, −7.231, and −7.114 respectively.

The selected top five compounds showed significant interactions with A42R protein, these interactions were aromatic and H-bonds, as detailed in the **(Figure 4** and **5**, **Table 2).** A42R-ExpressPick_1 formed two H-bonds with Glu3 and Arg127 residues and one arene bond with His124 residue. Whereas A42R-ExpressPick_2 interacted with Met1 residue though H-bond, A42R-ExpressPick_3 on the other hand displayed two H-bonds with Glu3, Arg127 residues, one ionic interaction with Glu3, and one arene bond with His124 residue. A42R-Emanime_1 demonstrated three H-bonds with Met1, His124, and Arg127 residues and one arene bond with Trp4 residue, while A42R-Natural Product_1 interacted through an H-bond with Trp4 residue of A42R.

**Figure 4:**
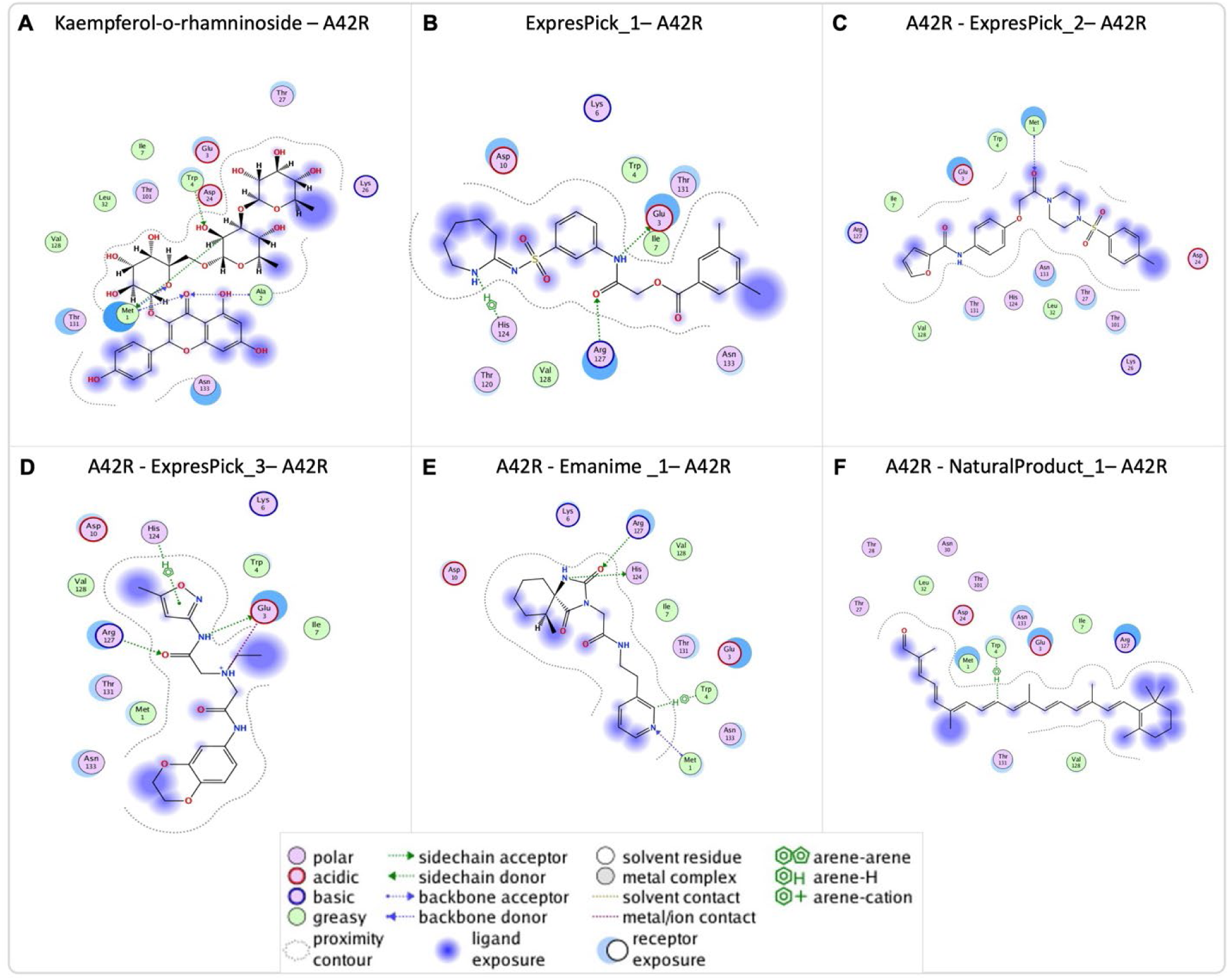
(A-D) The molecular interactions between the atoms of each ligand and the amino acid residues in the binding pocket of A42R are depicted in the 2D structure of the ligands. The sorts of interactions (ionic, hydrogen, and arene) as well as information on the polarity, acceptor/donor, ligand exposure, acidic/basic, and Greasy or neutral resides of the ligand are clarified.

**Figure 5:**
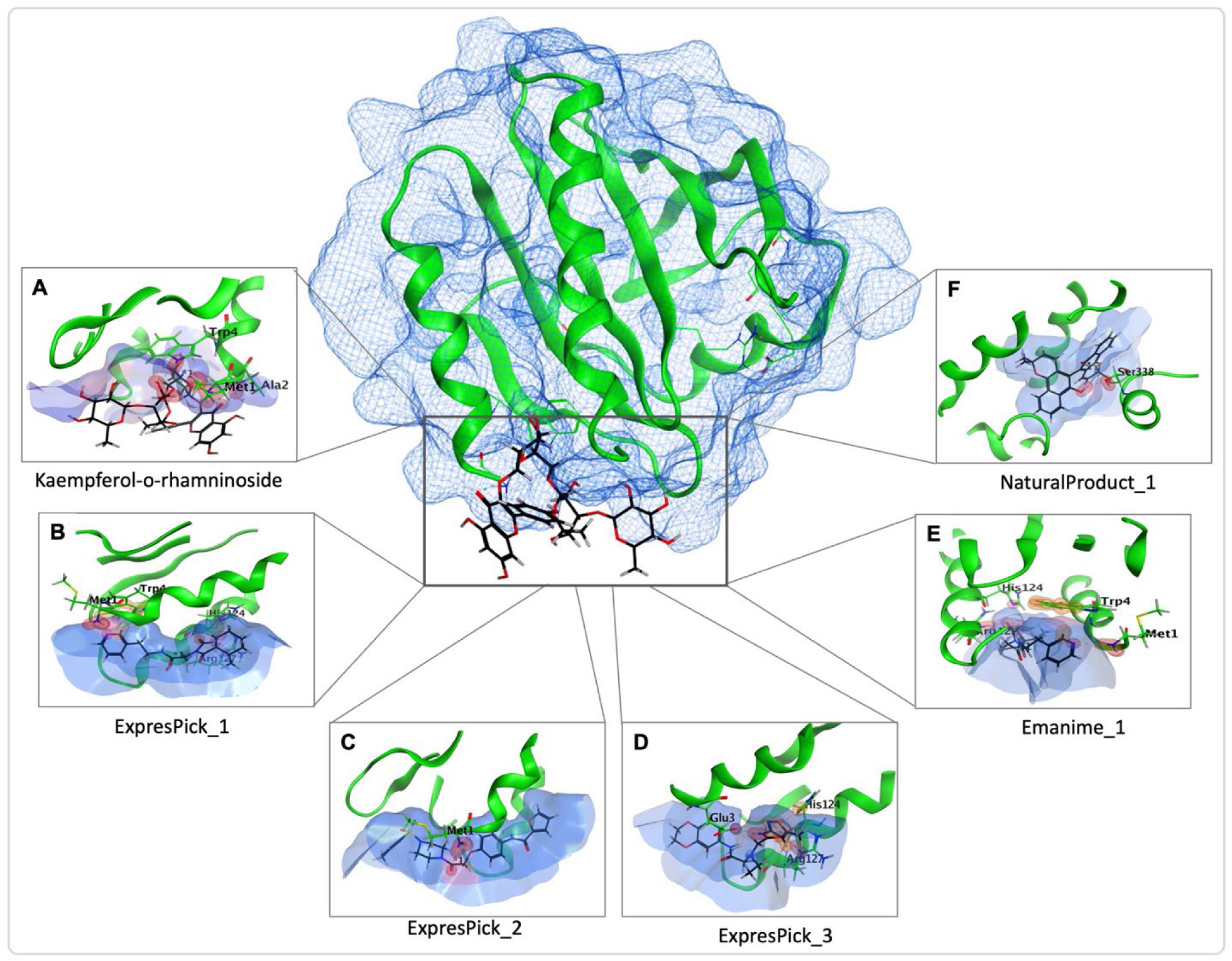
(A-D) Three-dimensional illustration of the ligand in the binding pocket of A42R, showing the atoms of the ligand, the corresponding protein residues that interact with it, and its orientation in three dimensions.

### Molecular dynamic simulations of DPol and A42R with selected ligands

Molecular dynamics (MD) simulations represent a widely utilized technique for assessing atomic behavior, structural stability, and atomic-level conformational changes ^31^. In this study, the top-ranking hits from the natural product library, MPD3 medicinal plant database, alkaloid library, flavonoid library, enamine antiviral library, and express-pick library underwent MD simulations using Schrodinger’s Desmond software (v. 2022-1).

#### Root Mean Square Deviation (RMSD)

The root mean square deviation (RMSD) serves as a prominent index in MD trajectory analysis, offering insights into the stability of conformations ^32^. Among the DPol-ligand complexes, DPol-ExpressPic_2, Gossypetin, and DPol-Natural Product_1, complexes exhibited the lowest RMSD, with values of 1.196 Å, 1.222 Å, and 1.245Å, respectively at 0.02 ns (**Figure 6A**). Over the course of the 1000 ns simulation, the DPol-Natural Product_1 and DPol-ExpressPic_2 complexes maintained almost similar RMSD, having average values of approximately 3.013Å and 3.031Å respectively. Notably, DPol-MPD3_1 and Gossypetin displayed consistent RMSD values throughout the simulations (average 2.229 and 2432 Å respectively). On average, the Gossypetin, DPol-MPD3_1, DPol-Natural Product_1, DPol-ExpressPic_1, DPol-Natural Product_2, and DPol-ExpressPic_2 complexes displayed RMSD values of 2.43Å, 2.23Å, 3.01Å, 3.93Å, 3.08Å, and 3.03Å, respectively.

**Figure 6:**
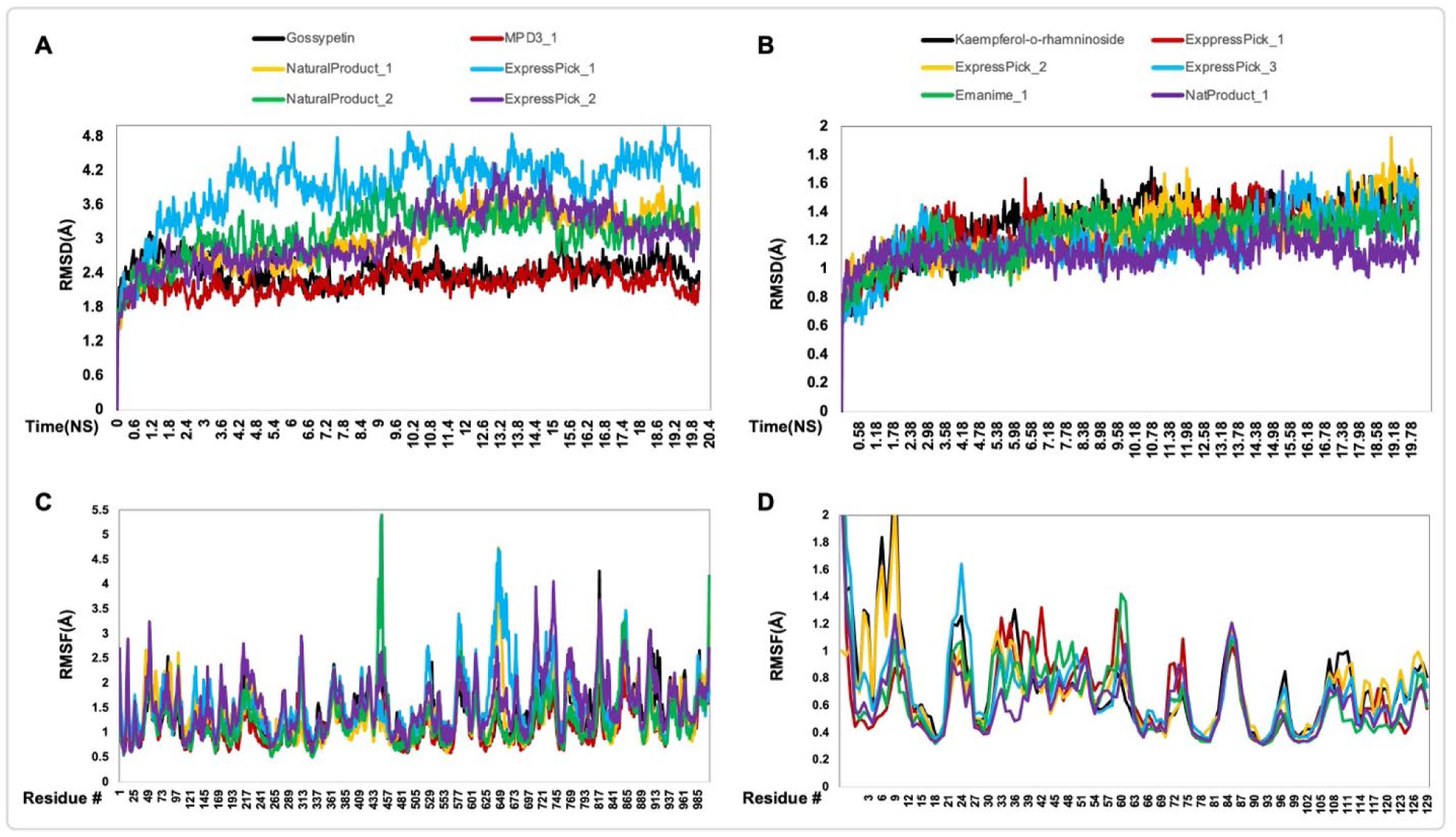
(A and B) Shown is the root mean square deviation (RMSD) values of the backbone and side-chain atoms of the hit compounds and the MPXV A42R and DPol complexes during the molecular dynamics (MD) simulation. (C and D) Illustration of the root mean square fluctuation (RMSF) values of the MPXV A42R and DPol proteins backbone and the hit compounds throughout the simulation.

However, among all A42R-ligand complexes, A42R-Emanime_1 and A42R-ExpressPick_2 complexes showed the lowest RMSD, with values of 0.581Å and 0.602Å, respectively at 0.02 ns **(Figure 6B).** Throughout the 1000 ns simulation, A42R-ExpressPick_3 and A42R-Emanime_1 complexes maintained almost similar RMSD, having average values of approximately 1.20Å and 1.23Å respectively. Whereas A42R-Natural Product_1 exhibited the lowest average RMSD (1.12Å). On average, the Kaempferol 3-O-rhamninoside, A42R-ExpressPick_1, A42R-ExpressPick_2, A42R-ExpressPick_3, A42R-Emanime_1, and A42R-Natural Product_1 complexes displayed RMSD values of 1.31Å, 1.28Å, 1.26Å, 1.20Å, 1.23Å, and 1.12Å respectively.

#### Root Mean Square Fluctuation (RMSF)

RMSF analysis allows for the observation of conformational shifts in residues during the simulation process ^4^. NatP-2 and NatP_1 in complexes with DPol exhibited the lowest RMSF, with values of 0.504 and 0.554, respectively, at residues 328 and 330 (**Figure 6C**). These two complexes maintained an average RMSF of 1.25 and 1.35, respectively. Conversely, DPol-Natural Product_2 and DPol-Natural Product_1 also exhibited the maximum fluctuations, with values of 5.41 and 4.75, respectively, at residues 446 and 645. Whereas the least average RMSF is maintained by DPol-MPD3_1 of value 1.14 throughout the simulation. On average, Gossypetin, DPol-MPD3_1, DPol-Natural Product_1, DPol-ExpressPic_1, DPol-Natural Product_2 and DPol-ExpressPic_2 complexes displayed mean RMSF values of 1.34, 1.14, 1.35, 1.54, 1.25, and 1.55, respectively.

While A42R-ExpressPick_2 and A42R-Emanime_1 on the other hand, showed the least RMSF values in complexes with A42R, having values 0.306 and 0.309 respectively at residue 95 **(Figure 6D).** Conversely, the A42R-ExpressPick_2 and A42R-ExpressPick_1 displayed the maximum RMSF of values 2.84 and 2.72, respectively at residue 1. However, the least average RMSF is exhibited by A42R-Natural Product_1 of value 0.62 throughout simulation. Overall, the average RMSF values of Kaempferol 3-O-rhamninoside, A42R-ExpressPick_1, A42R-ExpressPick_2, A42R-ExpressPick_3, A42R-Emanime_1, and A42R-NaturalProduct_1 complexes were 0.74, 0.68, 0.70, 0.71, 0.65, and 0.62 respectively.

Molecular dynamics simulations provide crucial insights into the behavior and stability of ligand-protein complexes, shedding light on their dynamic interactions and potential for therapeutic applications. The RMSD and RMSF analyses presented here offer valuable information on the stability and conformational dynamics of the studied complexes, with implications for drug discovery and development.

### Properties of selected ligands

The ligand RMSD indicates its items stability during the MD simulations. The RMSD monitored during MD simulations of 8HG1 and 4QWO proteins bound ligands revealed that Natural Product_2 (**Figure 7A**), and Enamine_1 and ExpressPick_3 (**Figure 7F**) shows better RMSD as compared to the reference ligands. The compactness of the complexes which primarily influences their rigidity was monitored using the radius of gyration. Higher radius of gyration values indicates greater instability in the complex, while lower values correspond to increased stability^33^. Throughout the simulation period, the radius of gyration values for Natural Product_2 shows almost similar values to that of reference (**Figure 7B**), on the other hand, enamine_1 and ExpressPick_3 shows better values as compared to the reference ligand (**Figure 7G**). A probe radius of 1.4 Å, equivalent to the van der Waals surface area of a water molecule, was used to calculate the molecular surface area (MolSA). Throughout the simulation, MolSA values exhibited fluctuations across different ranges. MPD3_1 and ExpressPick_2 and Kaempferol 3-O-rhamninoside show the higher MolSa values bound to DPol and A42R proteins of MPXV, respectively (**Figure 7C** and **H**). Solvent accessible surface area (SASA) values showed higher values for ExpressPick_1, ExpressPick_2 and Natural Product_2 shows better as compared to Gossypetin while Kaempferol 3-O-rhamninoside indicates better SASA scores (**Figure 7C** and **I**). The polar surface area (PSA) results better scores for both the reference ligands as compared to the selected test ligands (**Figure 7E** and **J**).

**Figure 7:**
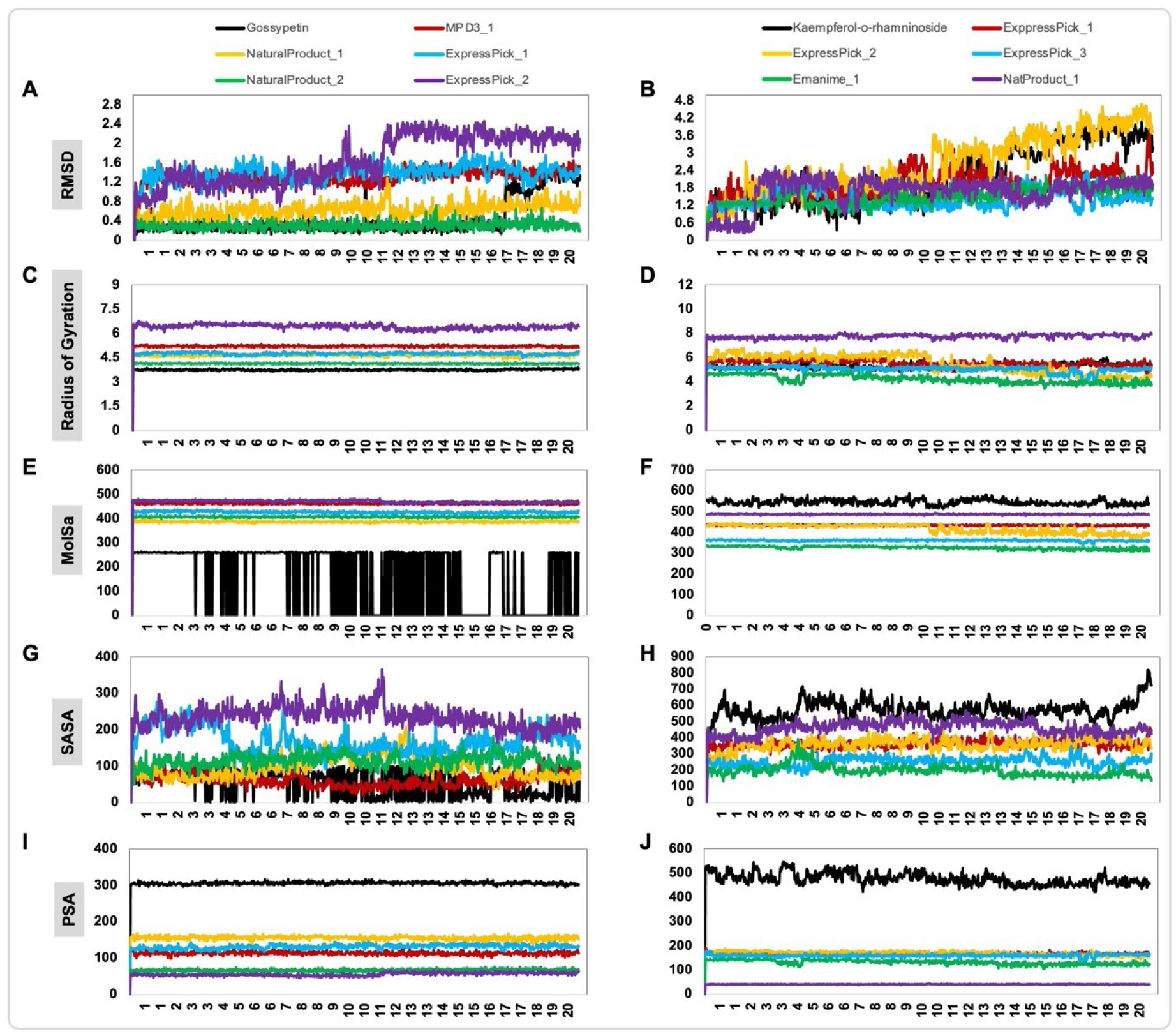
Properties of ligands during the MD simulations. (A and B) RMSD of ligands atoms, (C and D) radius of gyration, (E and F) molecular surface area (MolSA), (G and H) Solvent accessible surface area (SASA), (I and J) polar surface area (PSA).

### Pharmacokinetics and physiochemical properties of selected ligands

To distinguish between drug-like and non-drug-like compounds, Lipinski’s rule of five was utilized (**Table 3**). This rule helps identify a molecule’s drug-like properties based on its structural features^34,35^. Key pharmacokinetic parameters such as hydrogen bond acceptors, hydrogen bond donors, topological polar surface area (TPSA), bioavailability, molecular weight, and consensus log Po/w were assessed using MOE’s built-in descriptor compute tool.

**Table 3:** Thermotical ADMET properties of the selected ligands.

| Ligand-protein complex | Absorption |  |  | Distribution |  | Metabolism |  | Excretion |  | Toxicity |  |  |
| --- | --- | --- | --- | --- | --- | --- | --- | --- | --- | --- | --- | --- |
|  | Water solubility<br>Numeric (log mol/L) | Caco-2 permeability<br>Numeric (log Papp in 10-6<br>cm/s) | Human intestinal<br>absorption (%) | VDss (human) Numeric<br>(log mol/L) | BBB permeability Numeric<br>(log BBB) | CYP2D6 substrate (Y/N) | CYP12A inhibitor Y/N) | Total clearance<br>(log ml/min/kg) | Renal OCT2 substrate<br>(Y/N) | AMES toxicity (Y/N) | Skin sensitization (Y/N) | Hepatotoxicity (Y/N) |
| Gossypetin (Reference) -<br>DPol | -3.065 | -0.033 | 61.155 | 0.541 | -1.538 | N | N | -0.17 | N | N | N | N |
| MPD3_1- DPol | -3.991 | 1.245 | 94.505 | 0.679 | 0.015 | N | N | -0.411 | N | N | N | N |
| NaturalProduct_1- DPol | -3.736 | 0.647 | 0.647 | -0.282 | -1.154 | N | N | 0.513 | N | N | N | N |
| ExpressPic_1- DPol | -5.086 | 1.142 | 91.986 | -0.319 | -0.822 | N | N | 0.741 | N | Y | N | Y |
| Natural Product 2- DPol | -3.927 | 1.133 | 94.214 | -1.251 | -0.175 | N | N | 0.389 | N | N | N | Y |
| ExpressPic_2- DPol | -4.232 | 1.047 | 94.186 | 1.114 | 0.293 | Y | N | 0.675 | Y | N | N | N |
| Kaempferol 3-O-rhamninoside (Reference) -<br>A42R | -4.48 | -6.63 | 0.011 | 0.72 | -4.66 | Y | Y | 9.35 | N | N | N | N |
| ExpressPick_1 - A42R | -5.33 | 0.81 | 0.81 | -0.453 | -0.893 | N | N | -0.181 | N | N | N | N |
| ExpressPick_2 - A42R | -4.448 | 1.128 | 90.662 | -0.292 | -0.763 | N | N | 0.598 | Y | N | N | Y |
| ExpressPick_3 - A42R | -2.761 | 0.676 | 73.67 | -0.076 | -0.922 | N | N | 0.971 | N | N | N | Y |
| Emanime_1 - A42R | -3.654 | 0.221 | 67.602 | -0.25 | -0.876 | N | N | 1.337 | N | N | N | Y |
| Natural Product_1 - A42R | -8.113 | 1.361 | 0.318 | 0.318 | 0.784 | N | N | 1.368 |  | N |  | N |

Caco-2 permeability: the rate of flux of a compound across polarized Caco-2 cell monolayers, BBB: blood brain barrier, VDss: steady state volume of distribution, CYP2D6: Cytochrome P450 2D6, , CYP12A: Cytochrome P450 12A, OCT2: OCT2 is a primarily renal uptake transporter, AMES: carcinogenic effects of ligands on bacterial strain Salmonella typhimurium.

ADMET, representing Absorption, Distribution, Metabolism, Excretion, and Toxicity, encompasses crucial pharmacological properties for drug candidates. These properties are essential in drug development, as approximately 50% of drug failures are due to ADMET issues^36^. *In silico* ADMET analyses were conducted using the pkCSM web tool (https://biosig.lab.uq.edu.au/pkcsm/prediction)^37^. For selected ligands, various pharmacokinetic parameters such as water solubility, Caco-2 permeability, human intestinal absorption, blood-brain barrier (BBB) penetration, cytochrome P450 inhibition and substrate status, AMES toxicity, skin sensitization, and hepatotoxicity were evaluated.

As per Lipinski’s Rule of Five criteria in medicinal chemistry, compounds demonstrating druglikeness should possess a molecular weight (M. Weight) not exceeding 500, offer no more than 5 hydrogen bonds (lip_don), accept no more than 10 hydrogen bonds (lip_acc), and exhibit a partition coefficient (h_logP) less than 5 (**Table 4**)^38^. Non-compliance with these parameters may compromise the compound’s bioavailability. Nonetheless, in therapeutic development, deviations from these guidelines can be addressed through modifications in the administration route or by adding specific moieties to enhance the compound’s bioavailability.

**Table 4:**
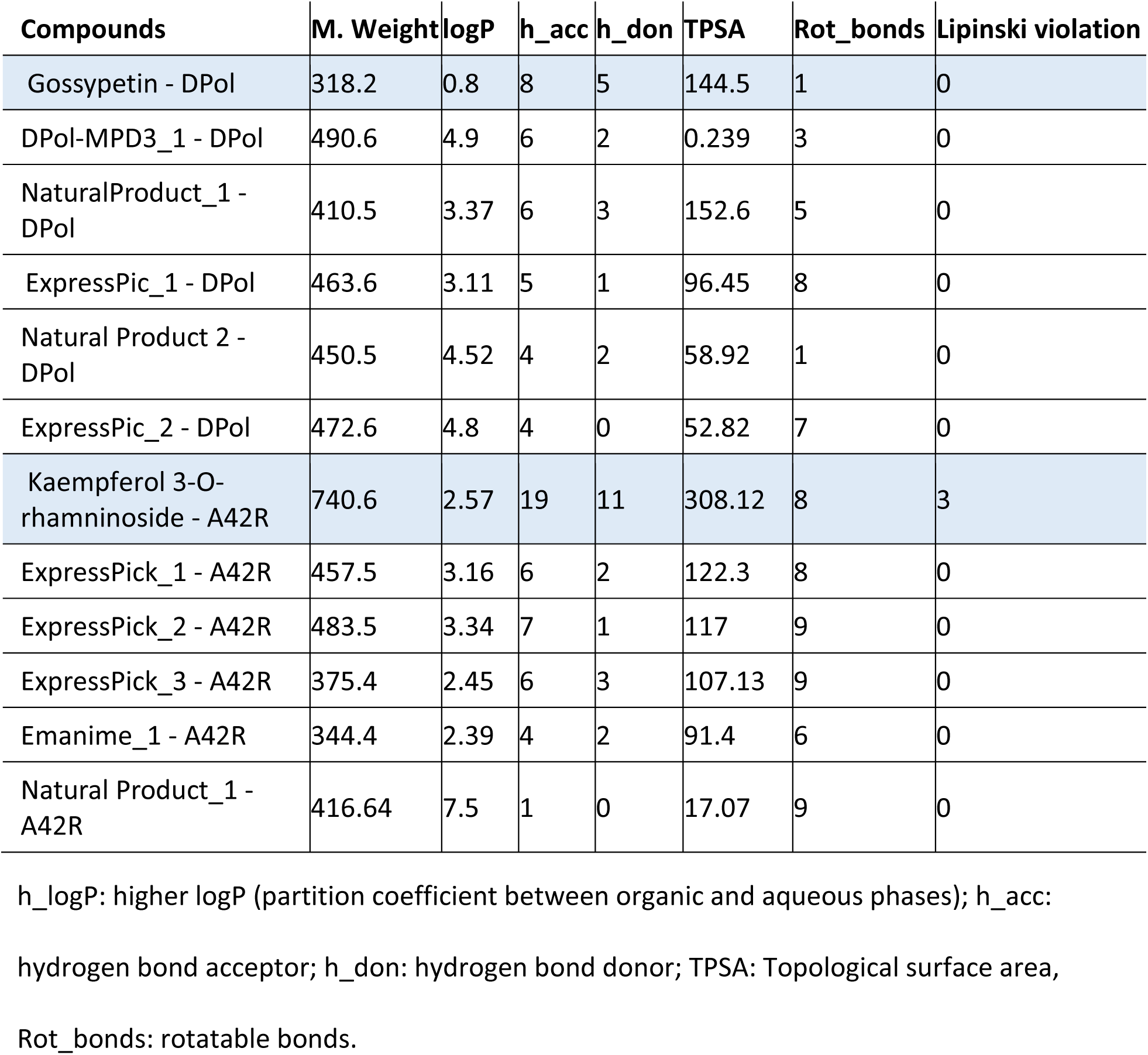
Physiochemical properties of the top 10 selected hit compounds from 5 libraries.

Furthermore, we checked the swissADME^29^ online database for oral bioavailibity of these compounds against the reference ligands (Gossypetin and Kaempferol 3-O-rhamninoside). Overall, most of the selected ligands shows better oral bioavailability than the already reported ligands (**Figure 8A-L**).

**Figure 8:**
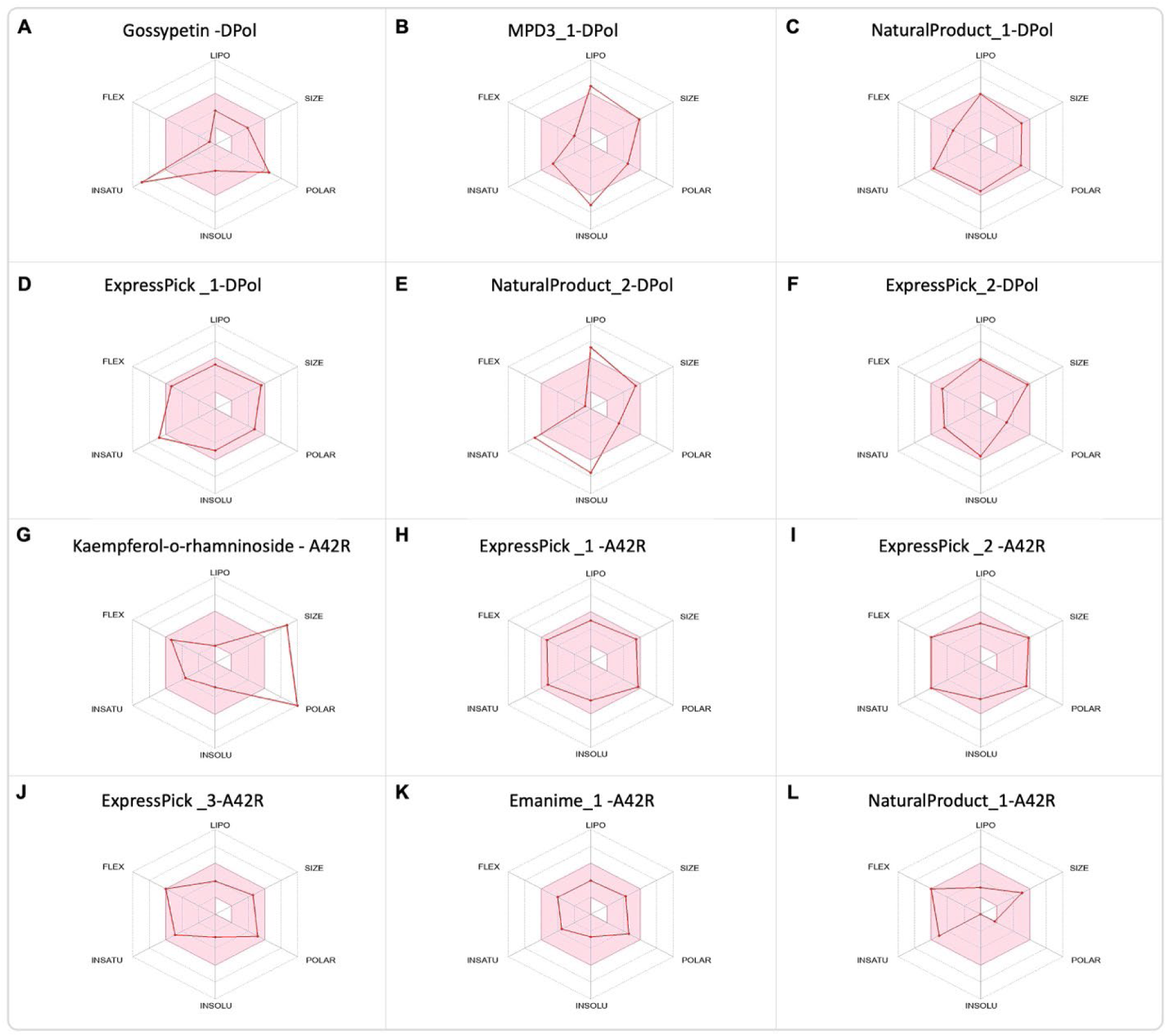
(A-L) The color zone represents the physiochemical space for oral bioavailability (retrieved from swissADME). LIPO (lipophilicity): −0.7<XLOGP3<+5.0; SIZE:<500g/mol; POLAR (Polarity): 20Å^2^ <TPSA<130Å^2^; INSOLU (Insolubility) −6<LogS (ESOL) < 0; INSATU (Instauration): 0.25 < fraction Csp3<1; FLEX (Flexibility): 0<Num. rotatable bonds <9.

Finally, we saved the selected ligands as sdf format using MOE and then converted it to Simplified Molecular Input Line Entry System (SMILES) using the online tool smiles translator and structure file generator^28^. Then the SMILES were used to retrieve the details of each ligands from PubChem^39^ such as their names, IDs, formulae and to confirm their molecular weight (**Table 5**).

**Table 5:**
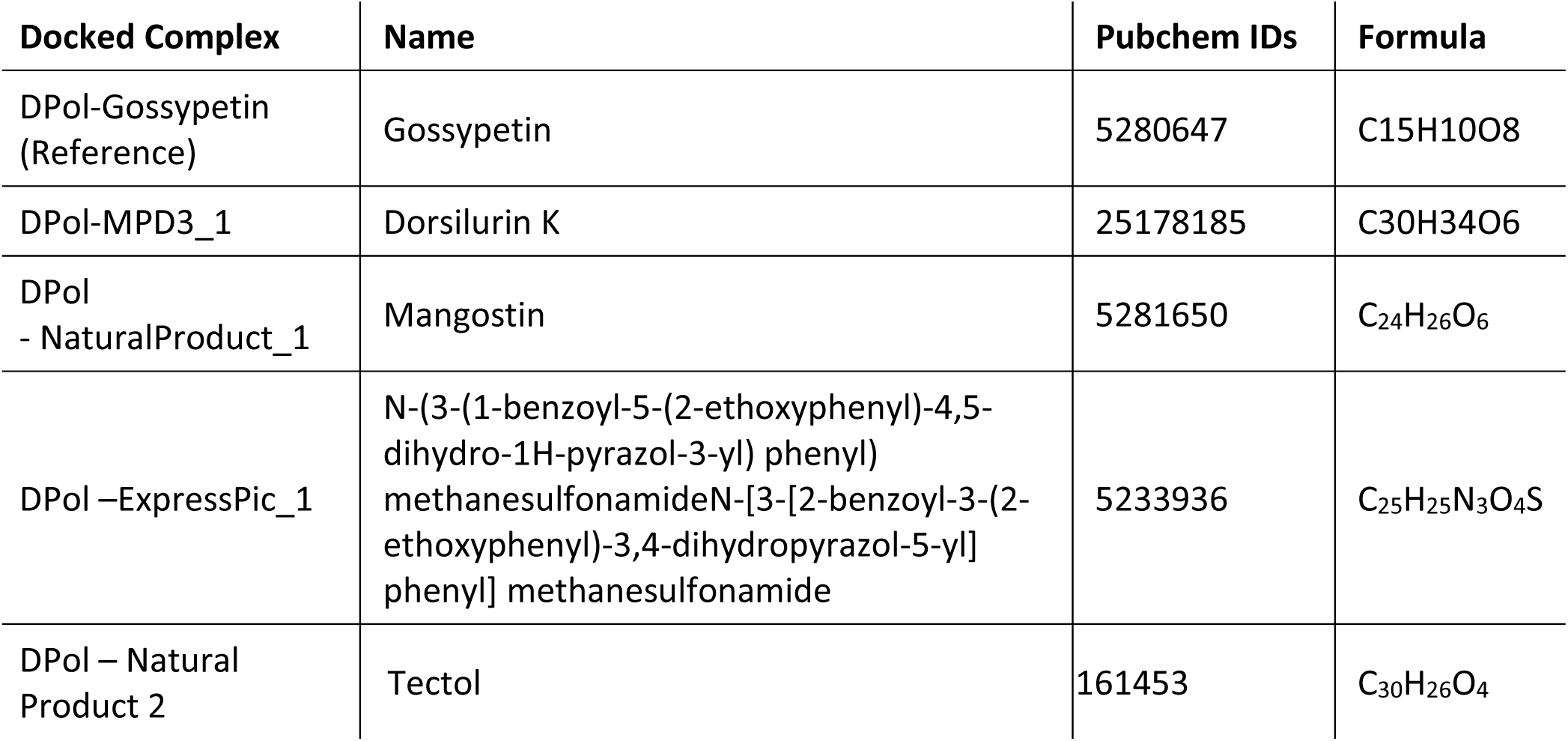
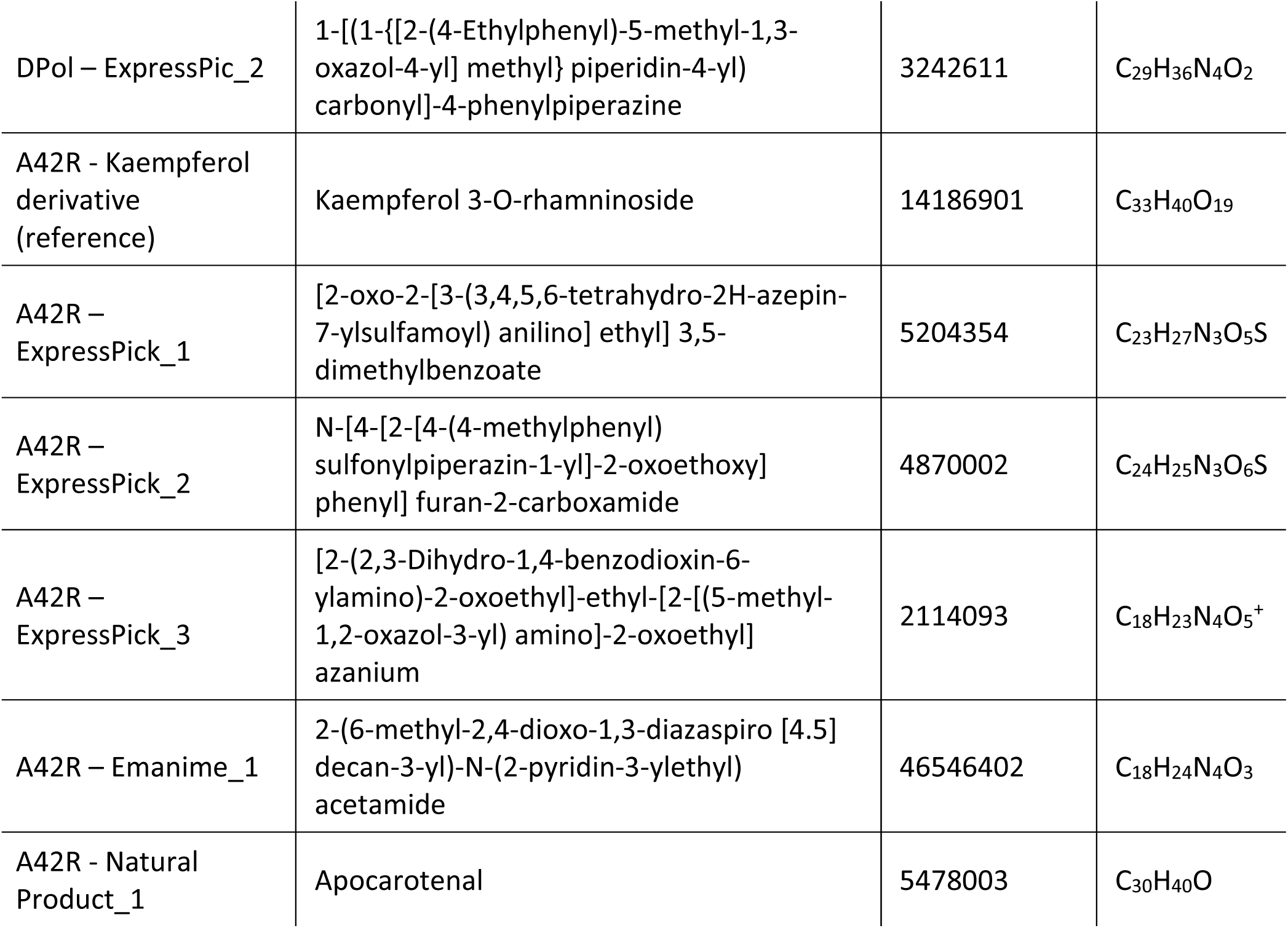
Docked complexes, their names, PubChem IDs and chemical formula.

| Docked Complex | Name | Pubchem IDs | Formula |
| --- | --- | --- | --- |
| DPol-Gossypetin (Reference) | Gossypetin | 5280647 | C<sub>15</sub>H<sub>10</sub>O<sub>8</sub> |
| DPol-MPD3_1 | Dorsilurin K | 25178185 | C<sub>30</sub>H<sub>34</sub>O<sub>6</sub> |
| DPol - NaturalProduct_1 | Mangostin | 5281650 | C<sub>24</sub>H<sub>26</sub>O<sub>6</sub> |
| DPol –ExpressPic_1 | N-(3-(1-benzoyl-5-(2-ethoxyphenyl)-4,5-dihydro-1H-pyrazol-3-yl) phenyl) methanesulfonamideN-[3-[2-benzoyl-3-(2-ethoxyphenyl)-3,4-dihydropyrazol-5-yl] phenyl] methanesulfonamide | 5233936 | C<sub>25</sub>H<sub>25</sub>N<sub>3</sub>O<sub>4</sub>S |
| DPol – Natural Product 2 | Tectol | 161453 | C<sub>30</sub>H<sub>26</sub>O<sub>4</sub> |
| DPol – ExpressPic_2 | 1-[(1-{[2-(4-Ethylphenyl)-5-methyl-1,3-oxazol-4-yl] methyl} piperidin-4-yl) carbonyl]-4-phenylpiperazine | 3242611 | C<sub>29</sub>H<sub>36</sub>N<sub>4</sub>O<sub>2</sub> |
| A42R - Kaempferol derivative (reference) | Kaempferol 3-O-rhamninoside | 14186901 | C<sub>33</sub>H<sub>40</sub>O<sub>19</sub> |
| A42R – ExpressPick_1 | [2-oxo-2-[3-(3,4,5,6-tetrahydro-2H-azepin-7-ylsulfamoyl) anilino] ethyl] 3,5-dimethylbenzoate | 5204354 | C<sub>23</sub>H<sub>27</sub>N<sub>3</sub>O<sub>5</sub>S |
| A42R – ExpressPick_2 | N-[4-[2-[4-(4-methylphenyl) sulfonylpiperazin-1-yl]-2-oxoethoxy] phenyl] furan-2-carboxamide | 4870002 | C<sub>24</sub>H<sub>25</sub>N<sub>3</sub>O<sub>6</sub>S |
| A42R – ExpressPick_3 | [2-(2,3-Dihydro-1,4-benzodioxin-6-ylamino)-2-oxoethyl]-ethyl-[2-[(5-methyl-1,2-oxazol-3-yl) amino]-2-oxoethyl] azanium | 2114093 | C<sub>18</sub>H<sub>23</sub>N<sub>4</sub>O<sub>5</sub><sup>+</sup> |
| A42R – Emanime_1 | 2-(6-methyl-2,4-dioxo-1,3-diazaspiro [4.5] decan-3-yl)-N-(2-pyridin-3-ylethyl) acetamide | 46546402 | C<sub>18</sub>H<sub>24</sub>N<sub>4</sub>O<sub>3</sub> |
| A42R - Natural Product_1 | Apocarotenal | 5478003 | C<sub>30</sub>H<sub>40</sub>O |

## DISCUSSION

Both humans and animals can contract monkeypox, a viral illness caused by the MPXV. MPXV requires a specialized DNA polymerase (DPol) enzyme for replication and dissemination within the host. This enzyme plays a pivotal role in manipulating the host’s immune response, contributing to the development of the monkeypox infection^40,41^. Consequently, comprehending the structure and function of DPol is of principal importance for the development of effective antiviral treatments and vaccines to combat monkeypox’s detrimental effects^41^.

While there are available treatments such as the JYNNEOS vaccine, tecovirimat, brincidofovir, and cidofovir for monkeypox, the exploration of adjuvant therapies utilizing phytochemicals and immune-enhancing diets holds promise in countering the spread of MPXV. Many phytochemicals have shown potential in reducing viral replication and bolstering host defenses, making them intriguing candidates for addressing viral diseases like influenza, HIV, herpes simplex virus, and SARS-CoV-2 ^42,43^. Although promising in preclinical research, the practical application of phytochemicals as antiviral medications faces challenges due to the cost of isolation and production, the need for extensive clinical investigations to determine optimal dosages and delivery methods, and the imperative for safety and efficacy ^43,44^.

In our study, we employed virtual screening techniques on various compound libraries, including the Natural product library, MPD3 medicinal plant database, Alkaloid library, Flavonoid library, Enamine antiviral library, and Express-pick library, to identify compounds capable of inhibiting DPol and A42R activities in MPXV. The docking analysis unveiled several compounds with notable interactions, including Dorsilurin K, Mangostin, N-(3-(1-benzoyl-5-(2-ethoxyphenyl)-4,5-dihydro-1H-pyrazol-3-yl)phenyl)methanesulfonamideN-[3-[2-benzoyl-3-(2-ethoxyphenyl)-3,4-dihydropyrazol-5-yl]phenyl]methanesulfonamide, Tectol, 1-[(1-{[2-(4-Ethylphenyl)-5-methyl-1,3-oxazol-4-yl]methyl}piperidin-4-yl)carbonyl]-4-phenylpiperazine, [2-oxo-2-[3-(3,4,5,6-tetrahydro-2H-azepin-7-ylsulfamoyl)anilino]ethyl] 3,5-dimethylbenzoate, N-[4-[2-[4-(4-methylphenyl)sulfonylpiperazin-1-yl]-2-oxoethoxy]phenyl]furan-2-carboxamide, [2-(2,3-Dihydro-1,4-benzodioxin-6-ylamino)-2-oxoethyl]-ethyl-[2-[(5-methyl-1,2-oxazol-3-yl)amino]-2-oxoethyl]azanium, 2-(6-methyl-2,4-dioxo-1,3-diazaspiro[4.5]decan-3-yl)-N-(2-pyridin-3-ylethyl)acetamide, and Apocarotenal, which exhibited impressive docking scores of −9.262, −8.634, −8.640, −8.426, −8.212, −7.703, −7.528, −7.382, −7.231, and −7.114 respectively (**Table 3**). Notably, these compounds displayed substantially lower docking scores compared to FDA-approved drugs for MPXV, namely, tecovirmat, bricindofovir, and cidofovir, which registered docking scores of −4.3, −7.5, and −4.5, respectively, in complexes with the modeled DPol of MPXV ^45^.

Furthermore, our molecular docking analysis identified Leu631, Arg634, and Lys661 as the key interacting residues within DPol, as they interacted with more than two ligands among the top selected ligands (**Figures 2** and **3**, **Table 2**). Leu631 formed H-bond with Gossypetin (reference), N-(3-(1-benzoyl-5-(2-ethoxyphenyl)-4,5-dihydro-1H-pyrazol-3-yl)phenyl)methanesulfonamideN-[3-[2-benzoyl-3-(2-ethoxyphenyl)-3,4-dihydropyrazol-5-yl]phenyl]methanesulfonamide and 1-[(1-{[2-(4-Ethylphenyl)-5-methyl-1,3-oxazol-4-yl]methyl}piperidin-4-yl)carbonyl]-4-phenylpiperazine. Like Leu631, Arg634 engaged in H-bonding interactions with Gossypetin (reference), N-(3-(1-benzoyl-5-(2-ethoxyphenyl)-4,5-dihydro-1H-pyrazol-3-yl)phenyl)methanesulfonamideN-[3-[2-benzoyl-3-(2-ethoxyphenyl)-3,4-dihydropyrazol-5-yl]phenyl]methanesulfonamide and 1-[(1-{[2-(4-Ethylphenyl)-5-methyl-1,3-oxazol-4-yl]methyl}piperidin-4-yl)carbonyl]-4-phenylpiperazine **(Figure 4** and **5**, **Table 2**). Whereas in A42R the most interacting residues were His124 and Trp4 which showed interactions with most of the selected ligands. His124 showed arene interactions with [2-oxo-2-[3-(3,4,5,6-tetrahydro-2H-azepin-7-ylsulfamoyl)anilino]ethyl] 3,5-dimethylbenzoate and [2-(2,3-Dihydro-1,4-benzodioxin-6-ylamino)-2-oxoethyl]-ethyl-[2-[(5-methyl-1,2-oxazol-3-yl)amino]-2-oxoethyl]azanium and H-bond 2-(6-methyl-2,4-dioxo-1,3-diazaspiro[4.5]decan-3-yl)-N-(2-pyridin-3-ylethyl)acetamide, while Trp4 displayed arene bond with 2-(6-methyl-2,4-dioxo-1,3-diazaspiro[4.5]decan-3-yl)-N-(2-pyridin-3-ylethyl)acetamide and Apocarotenal H-bond with Kaempferol 3-O-rhamninoside (reference ligand).

Subsequent molecular dynamics (MD) simulations highlighted the stability of the DPol and A42R complexes with various compounds. Among these complexes, 1-[(1-{[2-(4-Ethylphenyl)-5-methyl-1,3-oxazol-4-yl]methyl}piperidin-4-yl)carbonyl]-4-phenylpiperazine and Mangostin (with DPol) and 2-(6-methyl-2,4-dioxo-1,3-diazaspiro[4.5]decan-3-yl)-N-(2-pyridin-3-ylethyl)acetamide and N-[4-[2-[4-(4-methylphenyl)sulfonylpiperazin-1-yl]-2-oxoethoxy]phenyl]furan-2-carboxamide (with A42R) exhibited the lowest root mean square deviation (RMSD) values, ranging from 0.581Å to 1.245Å, affirming their structural stability (Figure 6A,B). The average RMSD values across all complexes(DPol and A42R with selected ligands), including Dorsilurin K, Mangostin, N-(3-(1-benzoyl-5-(2-ethoxyphenyl) −4,5-dihydro-1H-pyrazol-3-yl) phenyl) methanesulfonamide N-[3-[2-benzoyl-3-(2-ethoxyphenyl) −3,4-dihydropyrazol-5-yl] phenyl] methanesulfonamide, Tectol, 1-[(1-{[2-(4-Ethylphenyl) −5-methyl-1,3-oxazol-4-yl]methyl}piperidin-4-yl)carbonyl]-4-phenylpiperazine, Kaempferol 3-O-rhamninoside, [2-oxo-2-[3-(3,4,5,6-tetrahydro-2H-azepin-7-ylsulfamoyl)anilino]ethyl] 3,5-dimethylbenzoate, N-[4-[2-[4-(4-methylphenyl)sulfonylpiperazin-1-yl]-2-oxoethoxy]phenyl]furan-2-carboxamide, [2-(2,3-Dihydro-1,4-benzodioxin-6-ylamino)-2-oxoethyl]-ethyl-[2-[(5-methyl-1,2-oxazol-3-yl)amino]-2-oxoethyl]azanium, 2-(6-methyl-2,4-dioxo-1,3-diazaspiro[4.5]decan-3-yl)-N-(2-pyridin-3-ylethyl)acetamide, and Apocarotenal, remained ≤3.93 Å (**Figure 6**), further supporting the overall stability of these complexes ^46^.

Furthermore, we evaluated the ADMET, and physiological properties of our hit compounds and it revealed promising results as compares to the already reported ligands (**Figure 7** and **Table 4**). Finally, the oral bioavailability of these compounds also shows better results, in most cases except naturalproduct_2 ligand bound to DPol.

## CONCLUSION

The protein-ligand molecular docking results have unveiled ten noteworthy medicinal compounds: Dorsilurin K, Mangostin, N-(3-(1-benzoyl-5-(2-ethoxyphenyl)-4,5-dihydro-1H-pyrazol-3-yl)phenyl)methanesulfonamideN-[3-[2-benzoyl-3-(2-ethoxyphenyl)-3,4-dihydropyrazol-5-yl]phenyl]methanesulfonamide, Tectol, 1-[(1-{[2-(4-Ethylphenyl)-5-methyl-1,3-oxazol-4-yl]methyl}piperidin-4-yl)carbonyl]-4-phenylpiperazine, Kaempferol 3-O-rhamninoside, [2-oxo-2-[3-(3,4,5,6-tetrahydro-2H-azepin-7-ylsulfamoyl)anilino]ethyl] 3,5-dimethylbenzoate, N-[4-[2-[4-(4-methylphenyl)sulfonylpiperazin-1-yl]-2-oxoethoxy]phenyl]furan-2-carboxamide, [2-(2,3-Dihydro-1,4-benzodioxin-6-ylamino)-2-oxoethyl]-ethyl-[2-[(5-methyl-1,2-oxazol-3-yl)amino]-2-oxoethyl]azanium, 2-(6-methyl-2,4-dioxo-1,3-diazaspiro[4.5]decan-3-yl)-N-(2-pyridin-3-ylethyl)acetamide, and Apocarotenal, sourced from six distinct libraries, which demonstrate robust binding with the DPol and A42R proteins of MPXV. This molecular docking investigation has illuminated their high-affinity interactions with DPol and A42R, suggesting their potential to inhibit the replication of MPXV. Furthermore, the MD simulations have affirmed the stability of these lead compounds, exhibiting minimal deviations throughout the simulation period. Consequently, these compounds hold promise as prospective antiviral agents against MPXV. However, extensive preclinical research is imperative to ascertain their efficacy as antiviral medications.

## AUTHOR CONTRIBUTIONS

Muhammad Amjid and Tahir Muhammad conceptualized and designed the study, performed the *in-silico* experiments. Muhammad Amjid wrote the preliminary draft of the manuscript. Tahir Muhammad prepared the figures and helped with the writing of the manuscript. Muhammad Maroof khan helped in writing the initial draft of the manuscript, reviewed literature and helped in writing the preliminary draft of the manuscript. Stephen F. Pastore reviewed and edited the manuscript. John B. Vincent helped in the design of study reviewed and edited the manuscript.

Authors declare no competing interests

## DATA AVAILABILITY

The raw data supporting the findings of this study will be made available by the authors without undue reservation.

